# Krüppel-like factor 2- induced microRNAs: implications for treatment of pulmonary hypertension

**DOI:** 10.1101/583633

**Authors:** Hebah Sindi, Giusy Russomanno, Kyeong Beom Jo, Vahitha B. Abdul-Salam, Basma Qazi Chaudhry, Alexander J. Ainscough, Robert Szulcek, Harm Jan Bogaard, Claire C. Morgan, Soni Pullamsetti, Mai Alzaydi, Christopher J. Rhodes, Christina A. Eichstaedt, Ekkehard Grünig, Martin R. Wilkins, Beata Wojciak-Stothard

## Abstract

Flow-activated transcription factor Krüppel-like factor 2 (KLF2) signaling is compromised in pulmonary arterial hypertension (PAH). We aimed to identify KLF2-induced endothelium-protective exosomal microRNAs of potential therapeutic significance.

Eight exosomal microRNAs elevated by KLF2 but reduced in PAH were transfected into human pulmonary artery endothelial cells. Of these, only miR-181a-5p and miR-324-5p had anti-apoptotic, anti-inflammatory and anti-proliferative effect on endothelial cells and reduced proliferation of vascular smooth muscle cells *in vitro*. RNA sequencing of miRNA-transfected HPAECs revealed reduced expression of multiple genes implicated in vascular remodelling, including *ETS-1, NOTCH4, ACTA2, TNF-α, IL-1, MMP10, MAPK* and *NFATC2*. KLF2, miR-181a-5p and miR-324-5p were reduced, while their target genes were elevated in blood-derived endothelial colony forming cells and lung tissues from idiopathic and heritable PAH patients with disabling *KLF2* mutation and Sugen/hypoxia mice. Supplementation of miR-181a-5p and miR-324-5p or silencing of their target genes attenuated proliferative and angiogenic responses in endothelial cells from idiopathic PAH and prevented development of pulmonary hypertension in Sugen/hypoxia mice.

This study highlights potential therapeutic role of KLF2-induced exosomal microRNAs in PAH.

## INTRODUCTION

Pulmonary arterial hypertension (PAH) is a severe lung disorder characterised by progressive vascular remodelling and increased vasoconstriction. Endothelial damage followed by proliferation of vascular endothelial and smooth muscle cells underlie the disease pathology (Ranchoux, Harvey et al., 2018) and hypoxia and inflammation are known contributory factors (Rabinovitch, Guignabert et al., 2014). Formation of angio-obliterative vascular lesions, driven by vascular endothelial growth factor (VEGF), is a hallmark of severe PAH (Voelkel & Gomez-Arroyo, 2014).

Recently, a missense mutation in the transcription factor Krüppel-like factor 2 (*KLF2*) gene was identified in a family with autosomal heritable pulmonary arterial hypertension (HPAH), suggesting that KLF2 signalling may be compromised in the disease (Eichstaedt, Song et al., 2017).

KLF2 is activated by shear stress and plays a key role in the regulation of lung function and development (Anderson, Kern et al., 1995). KLF2-null mice exhibit abnormal blood vessel formation, resulting in embryonic haemorrhage and death. Within the blood vessel wall KLF2 is exclusively expressed in endothelial cells and promotes vascular homeostasis, counteracting inflammation, vascular leakage, thrombosis and VEGF-induced angiogenesis (Bhattacharya, Senbanerjee et al., 2005, SenBanerjee, Lin et al., 2004, Shi, Zhou et al., 2018). KLF2 also inhibits endothelial cell apoptosis, while promoting metabolic quiescence and reducing metabolic dependence on glucose (Doddaballapur, Michalik et al., 2015). The identification of missense mutation in *KLF2* gene is significant, as accumulating evidence implicates reduced KLF2 signalling in PAH pathogenesis. Inhibition of KLF2 expression correlates with increased PH severity in apelin knockout mice exposed to hypoxia (Chandra, Razavi et al., 2011). KLF2 overexpression improves pulmonary haemodynamics in hypoxic rats (Dungey, Deng et al., 2011), but can compromise liver function (Chen, Lu et al., 2014, Manavski, Abel et al., 2017), so other therapeutic approaches need to be identified. microRNAs (miRNAs) are small (∼22 nucleotide long) non-coding RNAs that negatively regulate gene expression at the posttranscriptional level (Small & Olson, 2011). Recent studies have shown that miRNAs released by the cells in exosomes, small membrane vesicles of 40–100 nm in diameter, can be taken up and modulate recipient cell responses in the immediate neighbourhood as well as in distant organs and tissues (Deng, Blanco et al., 2015, Zhang, Li et al., 2015).

Dysregulation of several miRNAs has been demonstrated in human and animal PAH but the choice of which miRNAs to target, constitutes a conceptual challenge (Negi & Chan, 2017). For example, the levels of KLF2-dependent miR-150 in plasma exosomes from PAH patients are reduced and correlate with survival (Rhodes, Wharton et al., 2013). A pioneering work by Hergenreider E *et al*. demonstrated that transfer of exosomal miRNAs from KLF2-overexpressing endothelial cells to underlying vascular SMCs reduces SMC de-differentiation, thus representing a promising strategy to combat atherosclerosis (Hergenreider, Heydt et al., 2012). Exosomes have gained special interest as carriers of miRNAs because of their transportability and the ability to convey information within the circulatory system (Zhang et al., 2015).

Our approach was to determine whether exosomal miRNAs from KLF2-overexpressing endothelial cells have vasculoprotective effects in PAH. We demonstrate dysregulation of KLF2-induced miRNA signalling in endothelial cells and lung tissues from idiopathic PAH (IPAH) patients and heritable PAH patients with *KLF2* mutation and present evidence of homeostatic and anti-remodelling effects of KLF2-induced miR-181a-5p and miR-324-5p *in vitro* and *in vivo*.

## RESULTS

### Endothelial exosomes mimic homeostatic effects of KLF2

In order to study the effects of KLF2-induced exosomes, KLF2 was overexpressed in human pulmonary artery endothelial cells (HPAECs) via adenoviral gene transfer. Recombinant KLF2 showed nuclear localisation (Figure 1A) and the level of KLF2 overexpression (∼3-fold increase) corresponded to expression level induced by physiological shear stress (10 dynes/cm^2^) in medium-size pulmonary arteries (Tang, Pickard et al., 2012).

**Figure 1.**
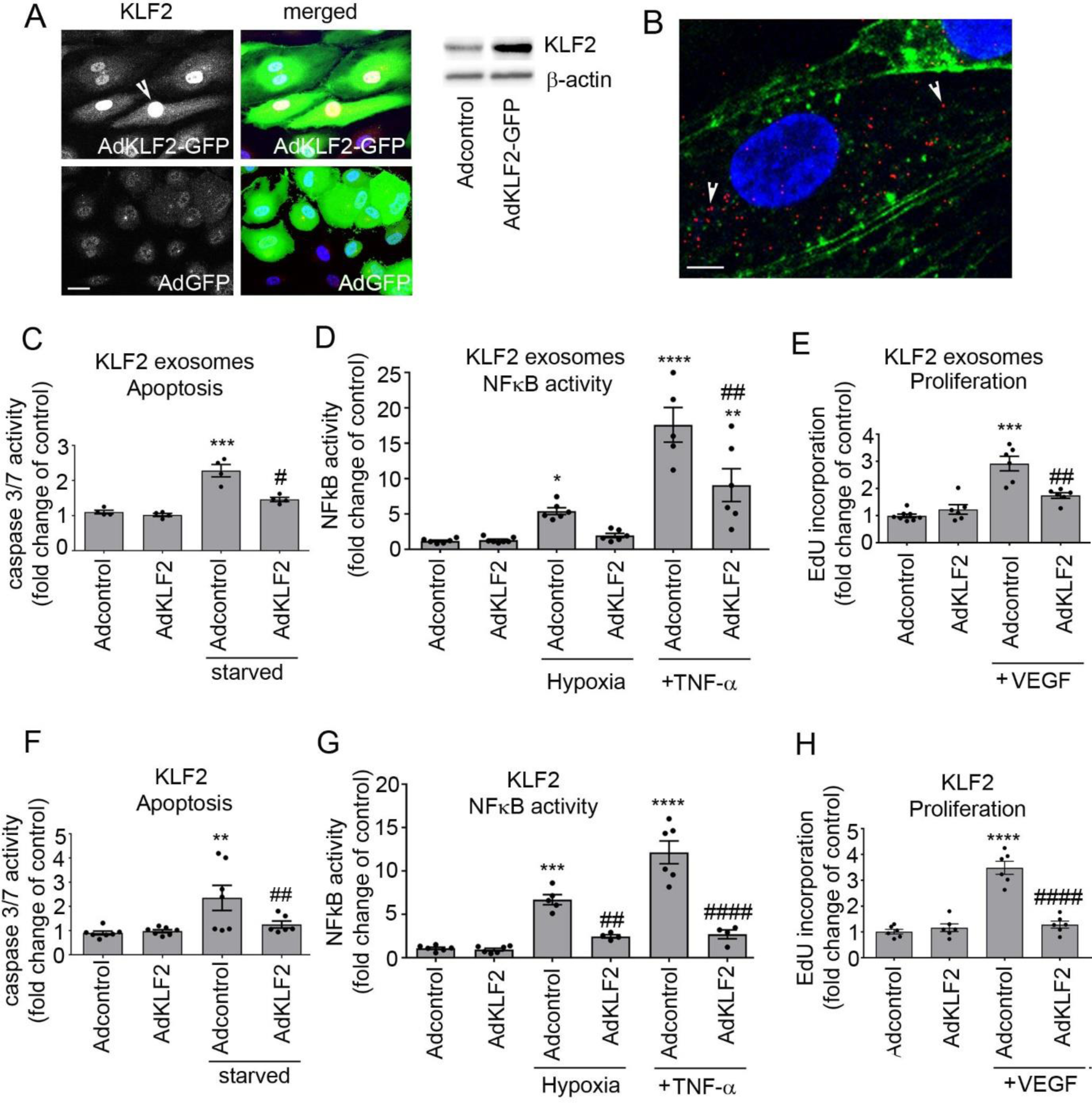
Effect of KLF2 and KLF2-induced exosomes on endothelial cell apoptosis, inflammatory activation and proliferation. (A) Expression of KLF2 in cells infected with AdGFP and AdKLF2-GFP (24h). Infected cells are green and the arrowhead points to nuclear localisation of KLF2; Bar=10µm. (B) Internalization of PKH26 Red-labelled exosomes (arrowheads) 1h after treatment. Bar=2µm. Effects of (C, D, E) KLF2-induced exosomes and KLF2 (F, G, H) on caspase 3/7 activation in serum-starved HPAECs (24h), hypoxia- and TNF-α)-induced (10 µg/L, 24h) NFκB activation and VEGF-induced (50ng/mL, 18h) cell proliferation in HPAECs, as indicated. Control exosomes were purified from AdGFP-expressing cells, while KLF2 exosomes were purified from AdKLF2-overexpressing HPAECs. **P<0.01; ***P<0.001, ****P<0.0001 comparison with Adcontrol. ^#^P<0.05, ^##^P<0.01, ^####^P<0.0001 comparison with treatment controls. One-way ANOVA with Tukey post-test. Values in (C-H) are mean fold-changes of Adcontrols ± SEM, n=4-7.

Exosomes were isolated from conditioned media collected from control (AdGFP) and AdKLF2-GFP-overexpressing (AdKLF2) HPAECs and the purity of the obtained fraction was confirmed by Nanosight LM10 particle tracking and exosomal protein marker analysis (Figure S1A-F in the Online Data Supplement). Conditioned media contained predominantly exosome-sized (<100 nm in diameter) particles, with approximately ∼2 x 10^10^ particles in each mL of medium. No significant differences in the total exosome number were found among the groups (Figure S1C, D in the Online Data Supplement). Purified exosomes were added to the cultured HPAECs at 1:1 donor-to-recipient cell ratio, ie. exosomes produced by 10 KLF2-overexpressing cells were added to 10 recipient cells in culture, corresponding to ∼10^5^ particles/cell. Internalization of fluorescently-labelled exosomes was evident after 1 hour of incubation (Figure 1B).

Treatment of HPAECs with KLF2 exosomes attenuated apoptosis and reduced hypoxia- and TNF-α-induced activation of the pro-inflammatory transcription factor NFκB (Figure 1C, D). KLF2 exosomes also inhibited VEGF-induced proliferation (Figure 1E and Figure S2 in the Online Data Supplement) and serum-induced cell proliferation (data not shown) in HPAECs, mimicking to a large extent the effects induced by KLF2 overexpression (Figure 1F-H).

### Identification of endothelium-protective exosomal miRNAs of potential therapeutic importance in PAH

miRNA profiling was carried out with miRCURY LNA™ Universal RT microRNA PCR (Human panel I+II) on exosome fractions from control (AdGFP) HPAECs and KLF2-overexpressing (AdKLF2) HPAECs. 330 miRNAs were detected per sample, with 183 miRNAs shared amongst all samples and 110 miRNAs differentially expressed, with a cut off p-value<0.05. Eighty-six of these miRNAs passed a Benjamini-Hochberg correction (p-adj<0.05) (Table S1 in the Online Data Supplement). Heat map with unsupervised hierarchical clustering performed on the 86 differentially expressed miRNAs is shown in Figure 2.

**Figure 2.**
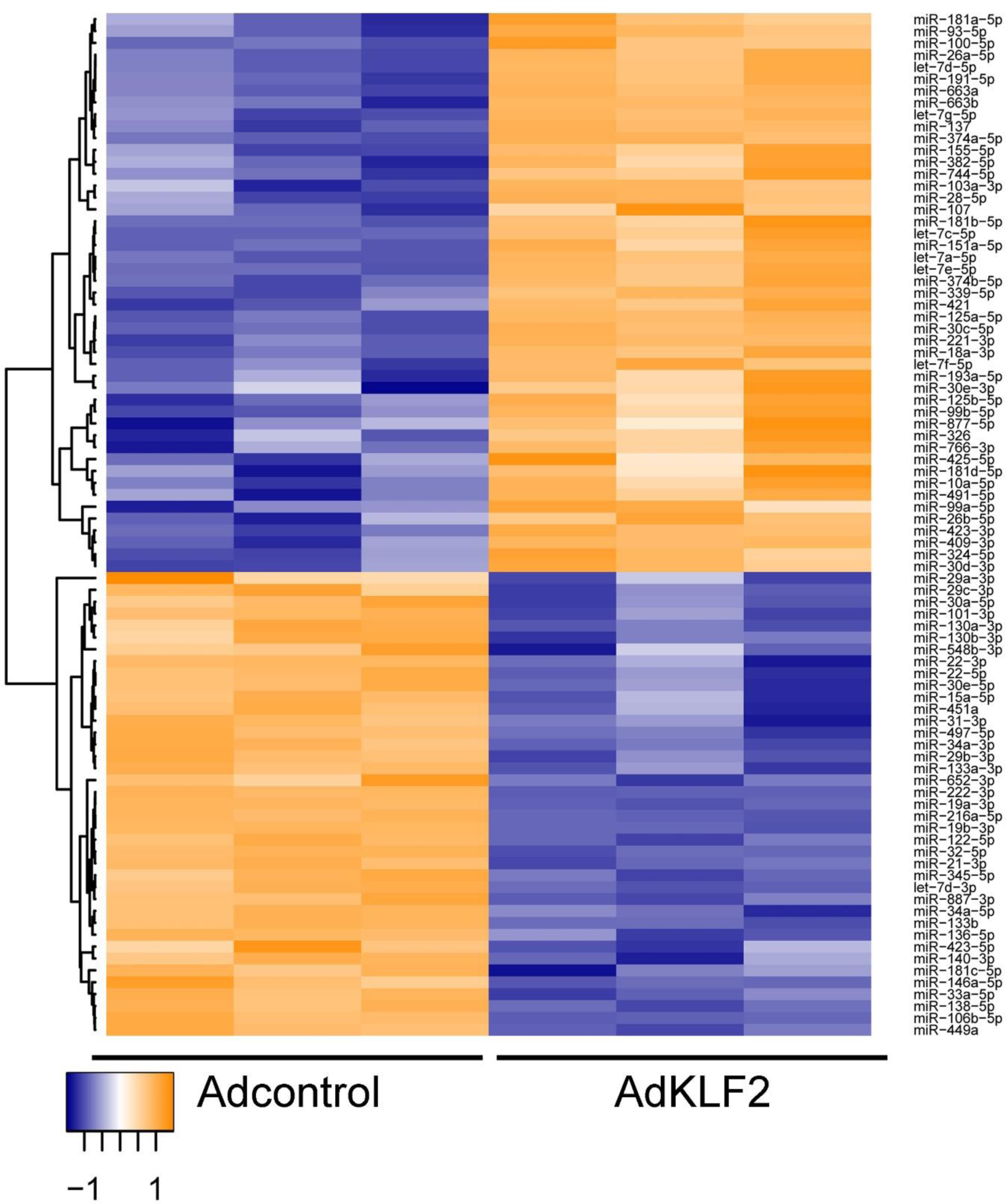
KLF2-induced changes in miRNA expression. (A) Heat map and unsupervised hierarchical clustering. The clustering was performed on 3 samples of exosomal pellets collected 24h post-infection with AdGFP or AdKLF2-GFP. Each row represents one miRNA, and each column represents one sample. The miRNA clustering tree is shown on the left. The colour scale illustrates the relative expression level of a miRNA across all samples: yellow colour represents an expression level above mean, purple colour represents expression lower than the mean.

The list of KLF2 miRNAs was then compared with published lists of miRNAs differentially expressed in human PAH and chronic hypoxia and MCT rat models of PAH (Caruso, MacLean et al., 2010, Rhodes et al., 2013). Eight miRNAs increased by KLF2 but reduced in human or animal PAH, were selected for further studies. The selected miRNAs comprised of let-7a-5p, miR-10a-5p, miR-125b-5p, miR-181a-5p, miR-191-5p, miR-30a-3p, miR-30c-5p, and miR-324-5p (Table S2 in the Online Data Supplement).

To verify whether miRNAs can pass from cell to cell under flow conditions, HPAECs were cultured in Ibidi flow chambers connected in tandem (Figure S3 in the Online Data Supplement). The cells grown in the first chamber were transfected with fluorescent Cy3-miR and then the unbound probe was washed away before the onset of flow at 4 dynes/cm^2^. The results show a directional transfer of fluorescent miRNA between endothelial cells under flow (Figure S3 in the Online Data Supplement).

### Endothelium-protective effects of miR-181 and miR-324

To verify the role of the 8 selected exosomal miRNAs, HPAECs were transfected with miRNA mimics prior to starvation, hypoxic exposure or stimulation of cells with TNF-α. Transfection efficiency, evaluated with Cy3-labelled miRNA, was ∼ 80 ± 5%.

Only miR-181a-5p and miR-324-5p (referred in the manuscript as miR-181 and miR-324), were protective in all study conditions (Figure S4 in the Online Data Supplement) and therefore were chosen for further analysis. Interestingly, while single treatments had only partial effect, the combined treatment with miR-181 and miR-324 was significantly more potent (Figure 3 A-D). Protective effects of miR-181/miR-324 treatment were prevented by specific miRNA inhibitors (Figure S5 in the Online Data Supplement).

**Figure 3.**
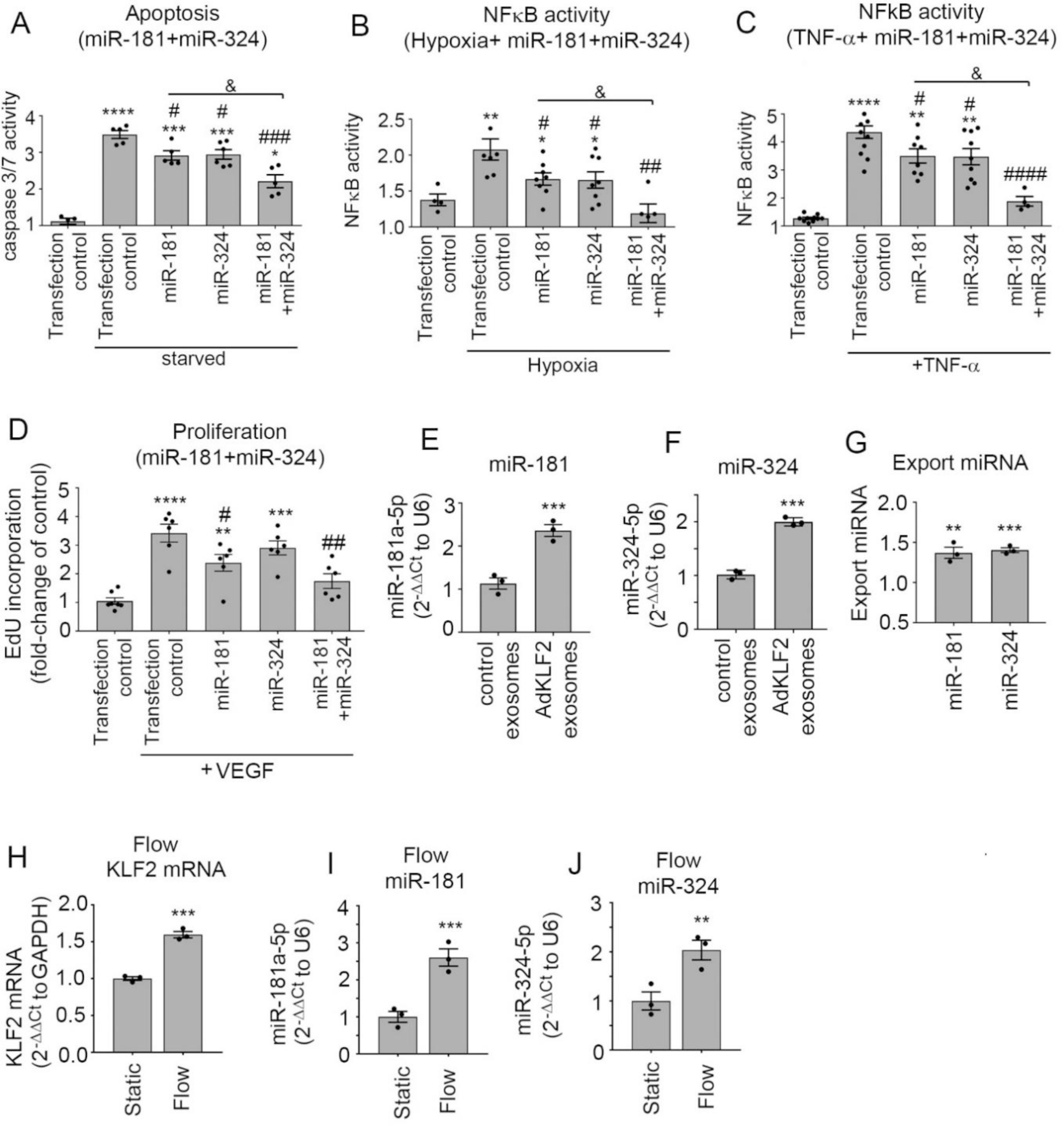
miR-181 and miR-324 mediate endothelium-protective effects of KLF2 exosomes. (A) Anti-apoptotic, (B, C) anti-inflammatory and (D) anti-proliferative effects of single and combined treatments with miR-181 and miR-324. HPAECs transfected with specific miRNAs were serum-starved or were infected with AdNFκB-luc and stimulated with hypoxia or TNF-α (10 µg/L) for 24h. Alternatively, HPAECc grown in serum-reduced medium were treated with VEGF (50ng/mL, 18h). (E) miR-181 and (F) miR-324 levels in HPAECs incubated with control or KLF2 exosomes for 24 h; qPCR. (G) Export ratios for miR-181and miR-324. (H) KLF2 mRNA, (I) miR-181 and (J) miR-324 changes in HPAECs under flow (10 dynes/cm^2^, 24h). Bars show mean fold-change of untreated controls ± SEM. *P<0.05, **P<0.01, ***P<0.001, ****P<0.0001, comparison with transfection controls; ^#^P<0.05, ^##^P<0.01, ^###^P<0.001,^####^P<0.0001, comparison with treatment controls or as indicated; ^&^P<0.05, comparison as indicated. T-test or one-way ANOVA with Tukey post-test, as appropriate. In (A-D) n=4-9, in (E-I) n=3.

miR-181 and miR-324 levels were significantly elevated in KLF2 exosomes, KLF2-overexpressing cells (Figure S6 in the Online Data Supplement) and in cells treated with KLF2 exosomes (Figure 3 E, F). The export ratio (the ratio between the exosomal and the intracellular levels of miRNA) was 1.37 for miR-181a-5p and 1.40 for miR-324-5p, indicating that these miRNAs are actively exported by the cells (Figure 3G). Expression levels of KLF2, miR-181 and miR-324 were also markedly elevated in flow-stimulated HPAECs (Figure 3 H-J).

To see if miR-181 and miR-324 play a role in endothelial-to-smooth muscle cell communication, human pulmonary artery smooth muscle cells (HPASMCs) were co-cultured with KLF2-overexpressing HPAECs in Transwell dishes. The two cell types were separated by a porous membrane (Figure 4A), which allows exchange of exosomal particles (Hergenreider et al., 2012). HPASMCs co-cultured with KLF2-overexpressing HPAECs showed elevated intracellular levels of miR-181 and miR-324 (Figure 4B, C) and reduced PDGF- and hypoxia-induced proliferation, compared with control cells (Figure 4D, E). The decrease in HPASMC proliferation was prevented by miR-181 and miR-324 inhibitors (Figure 4E).

**Figure 4.**
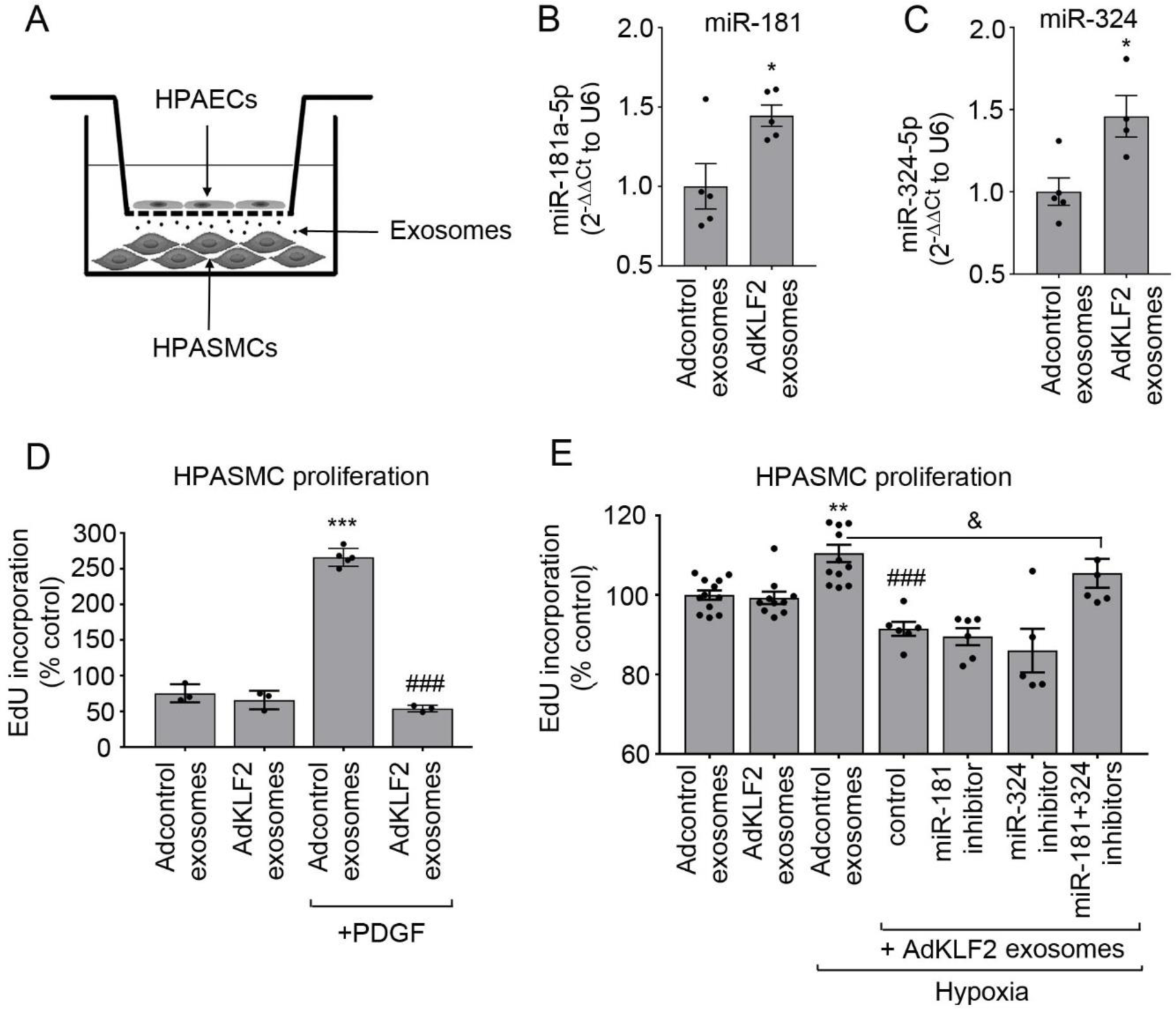
The effect of endothelial KLF2 on smooth muscle cell proliferation. (A) Transwell dish with KLF2-overexpressing HPAECs separated from HPASMCs by a porous (0.4 µm pore size) membrane. (B, C) miR-181 and miR-324 levels in HPASMCs co-cultured with Adcontrol and AdKLF2-overexpressing HPAECs for 24h. The effect of endothelial KLF2 on (D) PDGF (10 ng/mL, 48h)- and (E) hypoxia-induced HPASMC proliferation. In (D, E) bars show percentage of untreated, untransfected controls; *P<0.05, **P<0.01, ***P<0.001, comparison with Adcontrol/ exosomes group; ^###^P<0.001, comparison with Adcontrol + exosomes, hypoxia group; ^&^P<0.05, comparison as indicated. One-way ANOVA with Tukey post-test. Bars are means ± SEM, n=4-11.

### RNA profiling in miR-181 and miR-324-overexpressing HPAECs

HPAECs transfected with miR-181, miR-324 or non-targeting, control miRNA, were subjected to RNA profiling to identify targets of potential therapeutic significance.

The list of transcripts that were downregulated by miR-181 and miR-324 with fold-change >1.5 and adjusted p value <0.01 (977 for miR-181 and 930 for miR-324) and up-regulated (240 for miR-181 and 251 for miR-324) was compared with the list of predicted *in silico* target genes. 36 target genes of miR-181 and 37 target genes of miR-324 were then selected for pathway enrichment analysis (Figure 5A, Figure S7 and Tables S3, S4 in the Online Data Supplement). Gene Ontology (GO) and Kyoto Encyclopedia of Genes and Genomes (KEGG) pathway enrichment analysis of targets downregulated by miR-181 and miR-324 showed significant associations with TNF-α (p-adj=2.41×10^−6^), MAPK (p-adj=3.6×10^−4^), NFκB (p-adj=1.1×10^−4^), and Toll-like receptor signalling pathways (p-adj=1.2×10^−4^; VEGF (p-adj=1.32 × 10^−2^) (Figure 5B and Table S5 in the Online Data Supplement). Key targets of miR-181a-5p associated with inflammation, cell proliferation and vascular remodelling included α-SMA, TNF-α, IL-1, Notch4, MMP10, while targets of miR-324-5p included MAPK, NFATC2, ETS-1 (Table S5 in the Online Data Supplement). No significant KEGG pathway associations were found for targets up-regulated by miR-181 and miR-324.

**Figure 5.**
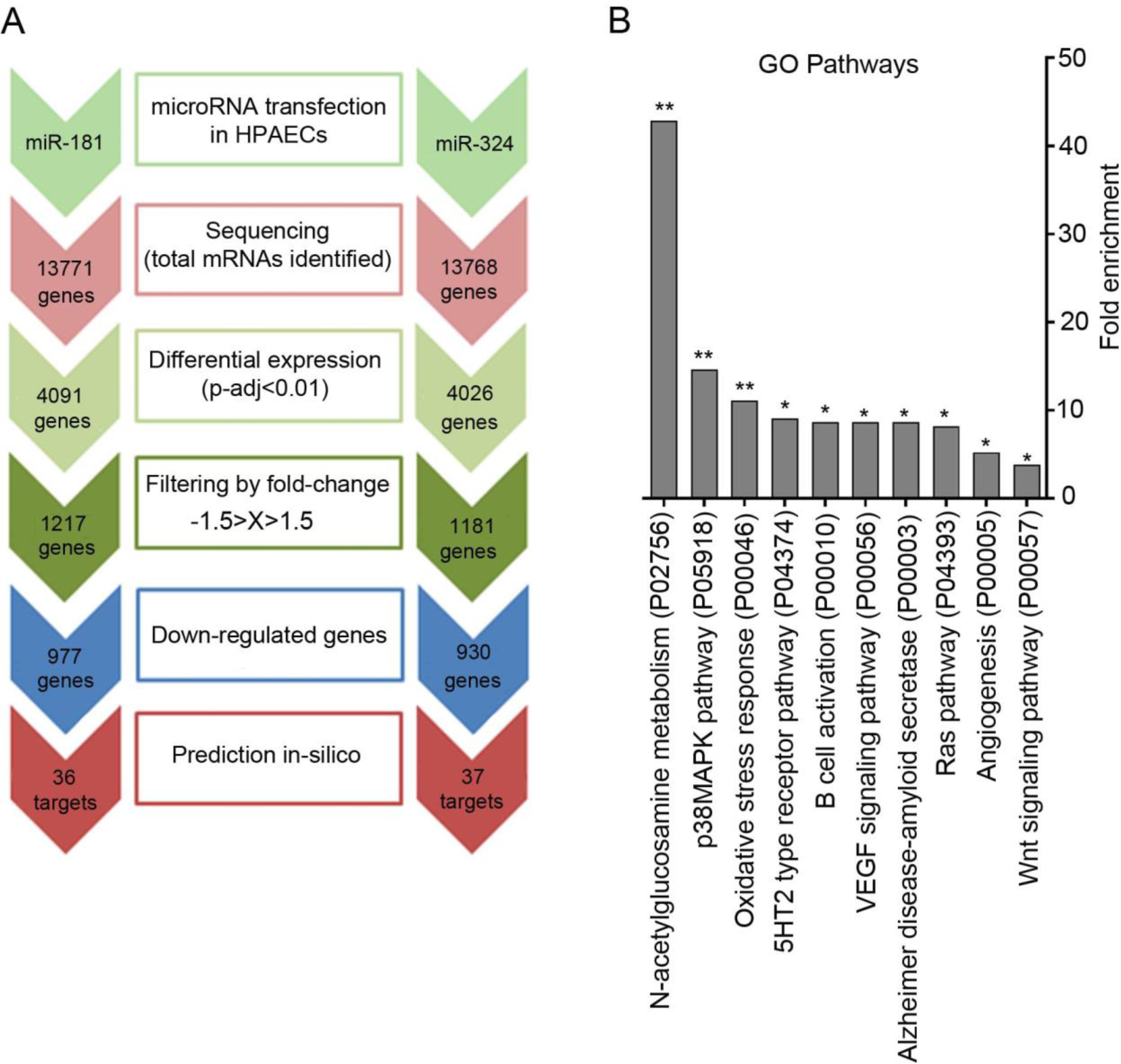
Identification of miR-181 and miR-324 targets using RNA sequencing and pathway analysis. (A) Flow chart shows identification of potential miRNA targets in PAH. HPAECs were transfected with miRNA mimics and subjected to RNA sequencing. miRNA target prediction was carried out with TargetScan Human, miRecords and Ingenuity Expert Findings (IPA). (B) Gene enrichment (GO pathways, Panther) in HPAECs transfected with miR-181 and miR-324. *p<0.05, **p<0.01

### Dysregulation of KLF2, miR-181 and miR-324 and their target genes in human PAH

Blood-derived endothelial colony-forming cells (ECFCs) are often used as surrogates for pulmonary endothelial cells in PAH. We examined the levels of KLF2, miR-181 and miR-324 and their selected gene targets in ECFCs from healthy volunteers (n=14) and IPAH patients (n=12). The IPAH cells showed a significant reduction in KLF2 mRNA, miR-181 and miR-324 and a marked increase in the levels of Notch4 (target of miR-181) and ETS-1 (target of miR-324) mRNAs, compared with healthy controls (Figure 6 A-E).

**Figure 6.**
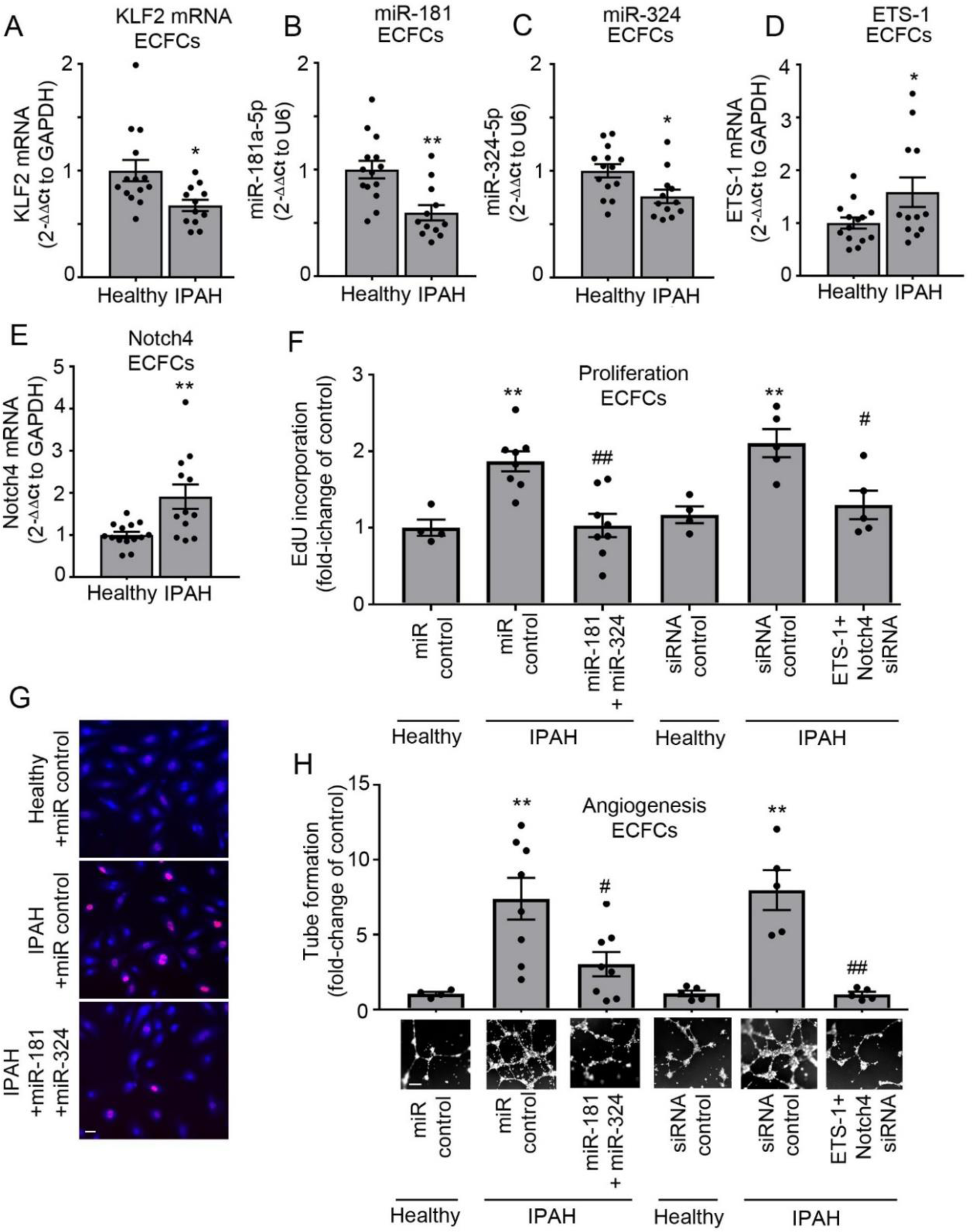
Dysregulation of KLF2 signalling in ECFCs from IPAH patients. (A) KLF2 mRNA, (B) miR-181 and (C) miR-324, (D) ETS-1 and (E) Notch4 expression in ECFCs from healthy volunteers (n=14) and IPAH patients (n=12). (F, G) Proliferation (EdU incorporation) and (H) angiogenesis (matrigel tube formation) in ECFCs from healthy individuals and IPAH patients, transfected with control miRNA, control siRNA, miR-181 and miR-324 or Notch4 and ETS-1 siRNA, as indicated. Representative corresponding images of tube formation in fluorescently-labelled live ECFCs are shown underneath the graph. In (G) EdU-incorporating nuclei are pink. In (G) Bar=20µm and in (H) Bar=100µm; *P<0.05, **P<0.001, comparison with corresponding controls; ^#^P<0.05, ^##^P<0.01, comparison between controls and miR-181/miR-324- or ETS-1/Notch4-treated IPAH ECFCs. Bars are means ± SEM, Student t-test, Mann-Whitney’s U test or one-way ANOVA with Tukey post-test, as appropriate.

Notch4 and ETS-1 are key regulators of endothelial proliferation and angiogenesis. Consistent with the elevated levels of Notch4 and ETS-1, IPAH ECFCs showed increased proliferation and tube formation in Matrigel, compared with healthy controls (Figure 6 F, G, H). These responses were markedly reduced in cells transfected with either miR-181/miR-324 or ETS-1 and Notch 4 siRNA (Figure 6 F, H). Reduction in ETS-1 and Notch4 expression in siRNA-transfected cells was confirmed by qPCR (Figure S8 in the Online Data Supplement).

The RNAscope fluorescent *in situ* hybridization, which allows specific identification and quantification of single transcripts (Wang, Flanagan et al., 2012), showed a marked reduction in the endothelial KLF2 mRNA and increased expression of Notch4 and ETS-1 mRNA in the remodelled vasculature of IPAH patients, compared with healthy lungs (n=6/group) (Figure 7 A-E).

**Figure 7.**
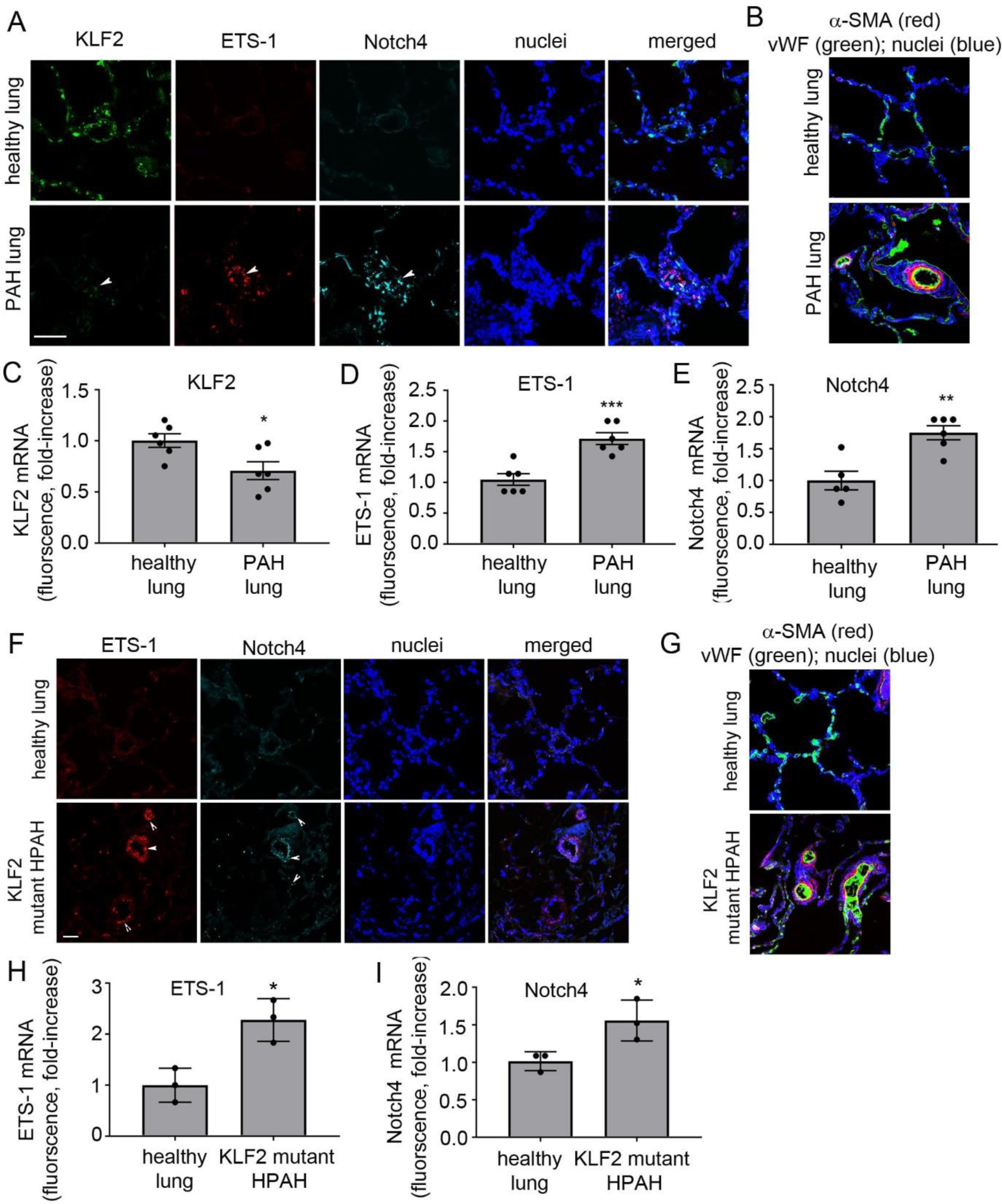
KLF2, ETS-1 and Notch4 mRNA levels in lung tissues of PAH patients. (A) Representative images of KLF2, ETS-1 and Notch4 mRNA distribution in healthy and PAH lung; RNAscope fluorescent *in situ* hybridization. Bar=50µm. (B) α-SMA and vWF staining in healthy and PAH lung. (C, D, E) Graphs showing KLF2, ETS-1 and Notch4 mRNA levels in healthy and PAH lung tissues; n=6. (F) Representative images of ETS-1 and Notch4 mRNA expression in healthy lung and lungs from HPAH patients with disabling *KLF2* mutation. Bar=25µm. (G) α-SMA and vWF staining in healthy and HPAH lung; (H, I) graphs showing ETS-1 and Notch4 levels in healthy lung tissues and lung tissues from HPAH patients with *KLF2* mutation, n=3. In (C, D, E, H, I) values are mean fold-changes of controls ± SEM *P<0.05, **P<0.01, ***P<0.001, comparison with healthy controls; Student t-test.

Lung tissues from HPAH patients with disabling c-terminal *KLF2* mutation (n=3) also showed a marked upregulation of Notch4 and ETS-1 in the remodelled vasculature, identified by endothelial vWF and a prominent α-SMA staining in the small, normally non-muscularised arterioles (Figure 7 F-I).

### miR-181 and miR-324 supplementation attenuates Sugen/hypoxia PH in mice

As a proof-of-concept, the Sugen/hypoxia mouse model of PAH was employed to study changes in the expression of KLF2 and miR-181 and miR-324 target genes in the remodelled lung, and to evaluate therapeutic potential of miR-181 and miR-324. In Sugen/hypoxia pre-clinical models of PAH, inhibition of VEGF receptor by Sugen (SU5416) induces an exaggerated vascular repair, driven largely by VEGF signaling, which further increases disease severity (Voelkel & Gomez-Arroyo, 2014).

Pilot experiments confirmed the enhanced effect of combined treatment (Figure S9 in the Online Data Supplement).

3-week exposure to Sugen/hypoxia induced a significant increase in the right ventricular systolic pressure (RVSP), right ventricular hypertrophy (RVH) and pulmonary vascular muscularisation in mice (Figure 8 A-C and H). Intravenous administration of miR-181 and miR-324 resulted in a significant increase in the levels of these miRNAs in the lungs of treated mice (Figure S9 in the Online Data Supplement).RVSP, RVH and vascular muscularization were significantly reduced in animals treated with miR-181/miR-324 (Figure 8 A-C, H).

**Figure 8.**
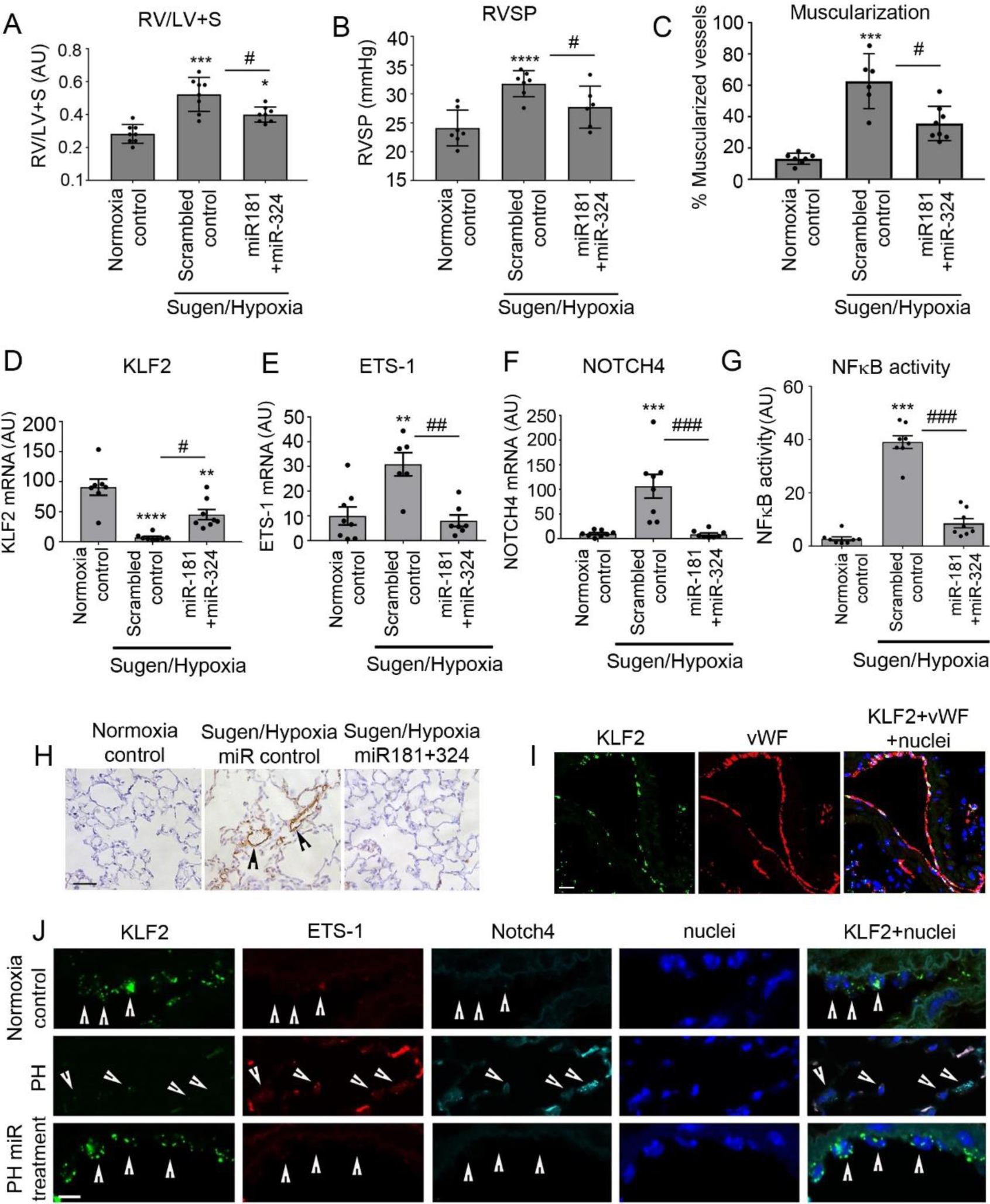
Delivery of miR181 and miR-324 attenuates pulmonary hypertension and reduces expression of Notch4, ETS-1 and NFκB activity in Sugen/hypoxia mice. (A) Right ventricular hypertrophy (RV/LV+S), (B) RVSP, and (C) vascular muscularisation in mice treated, as indicated. Changes in (D) KLF2, (E) ETS-1, (F) Notch4 mRNA levels and (G) NFκB activity (nuclear localisation) in the lungs of Sugen/hypoxia mice; fluorescence (arbitrary units). **P<0.01, ***P<0.001, ****P<0.0001, comparison with untreated controls; ^#^P<0.05, ^##^P<0.01, ^###^P<0.001, comparisons as indicated. Values are means ± SEM; one-way ANOVA with Tukey post-test, n=6-8. (H) Representative images of α-SMA staining in the untreated and miR-181+miR-324-treated mice lungs. (I) Co-localisation of KLF2 mRNA with vWF in mouse lung endothelium. (J) KLF2, ETS-1, Notch4 mRNA and nuclei in lung endothelial cells from healthy, pulmonary hypertensive (PH) and miRNA-treated mice, as indicated. Fluorescent *in situ* hybridization, confocal microscopy. Each fluorescent dot corresponds to a single transcript; arrowheads point to endothelial cells. In (H, I) Bar=25µm and in (J) Bar=10µm.

The levels of KLF2 mRNA in the lungs of PH mice were significantly lower compared with the lungs of healthy mice (Figure 8D). Changes in mouse body weight are shown in Figure S10 in the Online Data Supplement.

A marked reduction in KLF2 mRNA seen in the pulmonary vascular endothelium of Sugen/hypoxia mice was accompanied by increased expression of Notch4 and ETS-1 mRNA (Figure 8 E-F, I, J). Conversely, therapeutic supplementation of miR-181 and miR-324 resulted in a decrease in the mRNA and protein expression of Notch4 and ETS-1 and α-SMA, reduction in NFκB activity and reduction in the expression of cell proliferation marker, PCNA in the lung (Figure 8 E-G, J and Figures S11-S14 in the Online Data Supplement). Interestingly, endothelial KLF2 mRNA expression was partially restored in miRNA-treated mice, possibly as a result of improved flow conditions, secondary to the reduction in vascular remodelling (Figure 8D, J and Figure S11 in the Online Data Supplement). Endothelial localisation of the transcripts was confirmed by co-staining with vWF (Figure 8I).

## DISCUSSION

Exosome-mediated transfer of specific miRNAs between different cell types modulates vascular function (Aliotta, Pereira et al., 2016, Deng et al., 2015, Hergenreider et al., 2012). We sought to determine whether exosomal miRNAs from KLF2-overexpressing endothelial cells have a vasculo-protective effect in PAH. We show that delivery of selected KLF2-induced exosomal miRNAs improves pulmonary endothelial cell survival, reduces inflammatory responses and restricts vascular cell proliferation *in vitro* and *in vivo*.

To identify mediators of these effects, we conducted an unbiased screen of exosomal miRNAs and selected 8 miRNAs of interest, which were upregulated by KLF2 but reduced in PAH, with the view of potential therapeutic intervention. Of these, only miR-181 and miR-324 were protective in all experimental conditions and their effects were enhanced by the combined treatment. Functional convergence of multiple miRNAs on disease-relevant pathways has been demonstrated in the prevention of cardiac dysfunction and cancer (Chen, Duan et al., 2013, Huang, Tian et al., 2016, Negi & Chan, 2017, Su, Chang et al., 2013). miR-181 and miR-324 targets showed significant associations with key regulatory pathways in PAH, including TNF-α, MAPK, VEGF, NFκB and Toll-like receptor signalling. Numerous functional links among their respective, gene targets may explain, at least in part, their synergistic effects observed *in vitro* and *in vivo*.

The miR-181 family suppress NF-κB signalling and regulate endothelial cell activation, proliferation and immune cell homeostasis (Sun, Sit et al., 2014). Specifically, miR-181a-5p downregulates TNF-α, the key mediator of inflammatory responses in PAH (Hurst, Dunmore et al., 2017). In vascular SMCs miR-181a inhibits angiotensin II signalling, the adhesion of vascular SMCs to collagen and the expression of α-smooth muscle cell actin, suggesting that miR-181 may negatively modulate proliferative and migratory behaviour of HPASMCs in PAH (Remus, Lyle et al., 2013). We observed co-temporary, opposing changes in the expression of miR-181 and its target, Notch4, in the remodelled lungs of human PAH patients and Sugen/hypoxia mice. Notch4 predominantly localises to the vascular endothelium and regulates cell apoptosis, proliferation and VEGF-driven angiogenesis (Funahashi, Shawber et al., 2011, Hainaud, Contreres et al., 2006). Its expression is increased by VEGF and FGF (Marcelo, Goldie et al., 2013), both abundant in the remodelled PAH lung (Jonigk, Golpon et al., 2011, Voelkel & Gomez-Arroyo, 2014). Expression of Notch 1-4 is elevated in the remodelled lungs of hypoxic rats, with Notch4 mRNA showing the highest (5-fold) increase (Qiao, Xie et al., 2012). Consistent with this, we found that the levels of Notch4 mRNA were significantly elevated in blood-derived IPAH ECFCs, in the plexiform lesions in IPAH lungs (Jonigk et al., 2011) and in the remodelled lung tissues from HPAH patients with a *KLF2* mutation.

miR-324 controls proliferation, angiogenesis and oxygen metabolism in cells. We noted that miR-324 expression was significantly reduced, while expression of its key target, transcription factor ETS-1, was increased in ECFCs from IPAH patients, lung tissues from IPAH and HPAH patients and lungs from Sugen/hypoxia mice. ETS-1 activation is associated with expansive cell proliferation; high expression levels of ETS-1 facilitate the invasive behaviour of tumour cells (Zhou, Zhou et al., 2018) and are predictive of the poor prognosis in multiple types of carcinoma (Dittmer, 2015). ETS-1 also regulates VEGF-induces transcriptional responses (Chen, Fu et al., 2017), increases endothelial angiogenesis, extracellular matrix remodelling and the expression of glycolytic enzymes and associated feeder pathways (Verschoor, Wilson et al., 2010), known to augment vascular remodelling in PAH (Cao, Xie et al., 2015, Ryan & Archer, 2015).

ECFCs from IPAH patients display many of cellular abnormalities associated with the disease, including hyperproliferation, mitochondrial dysfunction and impaired ability to form vascular networks, reflective of the exaggerated repair processes in the remodelled lung (Caruso, Dunmore et al., 2017, Duong, Erzurum et al., 2011, Ormiston, Toshner et al., 2015, Smits, Tasev et al., 2018, Wojciak-Stothard, Abdul-Salam et al., 2014a). We observed that proliferative and angiogenic responses of IPAH ECFCs were reduced by overexpression of miR-181 and miR-324 and by silencing of Notch4 and ETS-1.

Homeostatic effects of KLF2 exosomes are likely to result from the cumulative reduction in expression of multiple targets of miR-181 and miR-324. Considering that miRNAs regulate the expression levels of multiple genes simultaneously and often cooperatively (Small & Olson, 2011), delineating individual contributions of miR-181 and miR-324 gene targets is beyond the scope of this investigation.

KLF2 encourages selective sorting of miRNAs into exosomes (Jae, McEwan et al., 2015) and our data indicate that miR-181a-5p and miR-324-5p are actively exported by the cells. Although the mechanism of miRNA sorting is not well understood (Zhang et al., 2015), specific consensus sequence motifs (EXOmotifs) have been implicated (Villarroya-Beltri, Gutierrez-Vazquez et al., 2013). miR-181a-5p has two EXOmotifs that differ from consensus sequence in one nucleotide (GGAC, UGAC), while miR-324-5p shows three motifs, one with perfect match to consensus sequence (CCCU) and two showing alteration in one nucleotide (UCCG, GGAC).

Mechanisms responsible for the reduction in KLF2 expression and signalling in PAH are currently unknown but may involve dysregulation of BMPR2 function (Eichstaedt et al., 2017) or inflammation (Kumar, Lin et al., 2005). Inhibition of KLF2 signalling is likely to result in maladaptive responses to flow and endothelial damage (Bonnet & Provencher, 2016). Interestingly, endothelial cells isolated from the subpleural lung microcirculation of patients with PAH at the time of transplantation or necropsy and submitted to *ex vivo* high shear stress exhibit delayed morphological adaptation, compared with controls (Szulcek, Happe et al., 2016).

In summary, we show that homeostatic effects attributed to KLF2 (Bhattacharya et al., 2005) and laminar shear stress (Doddaballapur et al., 2015) can be mimicked, at least in part, by KLF2-induced exosomal miRNAs. This study provides evidence of dysregulated KLF2 signaling in PAH and highlights the potential therapeutic role of KLF2-regulated exosomal miRNAs in pulmonary hypertension and other diseases associated with endothelial damage, inflammation, proliferation and propensity for vascular cell enlargement.

## MATERIALS AND METHODS

A full description of materials and methods is provided in the Online Data Supplement. Accession numbers for miRNA microarray and RNA sequencing data will be provided upon provisional acceptance of the manuscript.

### Cell culture

Human pulmonary artery endothelial cell (HPAECs, Promocell, C-12241) and human pulmonary artery smooth muscle cells (HPASMCs, Lonza, CC-2581) were cultured in endothelial cell growth medium 2 (ECGM2; PromoCell, C-22111), and smooth muscle cell growth medium 2 (SMCGM2, PromoCell, C-22062), as previously described (Wojciak-Stothard et al., 2014a). In some experiments, the cells were incubated with human recombinant TNF-α (R&D, 210-TA-020; 10 µg/L) or exposed to hypoxia (5% CO_2_, 2%O_2_) for 24-72 hours.

For non-contact co-culture of HPAECs and HPASMCs, Transwell dishes with 0.4 μm pore polyester membrane inserts (Scientific Laboratory Supplies, UK) were used.

### Blood-derived human endothelial cell culture

Human endothelial colony forming cells (ECFCs), were derived from peripheral blood samples as previously described (Wojciak-Stothard et al., 2014a). Clinical information is shown in Table S6 in the Online Data Supplement).

### Adenoviral gene transfer

HPAECs (80-90% confluent) were infected with adenoviruses adeno-GFP (Adcontrol; Vector Biolabs, 1060) or adeno-KLF2-GFP (AdKLF2; Vector Biolabs, ADV-213187) at the multiplicity of infection (MOI) 1: 100 or 1:250 and used for experiments 24h post-infection.

To stain KLF2 and F-actin in cells, fixed and permeabilized HPAECs were incubated with mouse monoclonal anti-KLF-2 antibody (Novus Biologicals, NB100-1051; 1:1000) and then FITC-Goat Anti-Mouse IgG (Jackson ImmunoResearch Inc., 115-095-003; 1:200) and F-actin stain, 1mg/L TRITC-phalloidin (Sigma, P1951). Cells were examined under the fluorescent confocal microscope (Leica, TCS SP5, Leica Biosystems).

### Exosome purification and quantification

Exosomes were isolated from conditioned media of HPAECs overexpressing AdGFP or AdKLF2-GFP cultured in ECGM2 medium (PromoCell, C-22111) supplemented with 2% exosome-depleted foetal bovine serum (Thermo Fisher, A2720801) using miRCURY™ Exosome Isolation kit (Exiqon; 300102). The particle size distribution and number in conditioned media and purified exosome fractions were studied with NanoSight LM10 Particle Size Analyzer and Particle Counter (Malvern Instruments Ltd). Exosome marker proteins were analysed by western blotting. The purity of exosome fraction was also evaluated with Exosome Array (System Biosciences, EXORAY-4).

### Cell treatment with exosomes and imaging of exosome internalization

Exosomes purified from 10 ml of conditioned medium were re-suspended in 10 ml of fresh ECGM2 medium and added to HPAECs grown in 96- or 24-well dishes. To visualise the internalised exosomes, exosomal cell membrane was stained with PKH26 Red Fluorescent Cell Membrane Linker Kit (Sigma-Aldrich; PKH26GL-1KT), according to the manufacturer’s protocol. Labelled exosomes were incubated with endothelial cells for 1h and then the cells were fixed, permeabilised, stained with 1μg/ml FITC-phalloidin, washed and mounted in Vectashield containing nuclear stain DAPI. Images were taken under the Leica TCS SP5 Confocal Microscope.

### Universal RT microRNA PCR Human panel I+II

miRNA profile in HPAECs overexpressing AdGFP (controls) and AdKLF2-GFP was analysed with Exiqon miRCURY LNA™ Universal RT (microRNA PCR Human panel I+II). The list of KLF2-induced miRNAs was then compared with the published lists of differentially expressed miRNAs in PAH patients and PH animals (Caruso et al., 2010, Rhodes et al., 2013). miRNAs that were reduced in PAH but elevated by KLF2, were selected for further analysis.

### Cell Transfection

Transfection of miRNA mimics and inhibitors was carried out with Lipofectamine RNAiMAX Transfection Reagent (Thermo Fisher, 13778150). Briefly, HPAECs were left untreated (control) or were transfected with control miRNA (non-targeting transfection control; Ambion Life Technologies, 4464076) at 20 nmol/L or control miRNA at 10 nmol/L plus either miR-181a-5p (10 nmol/L; Ambion Life Technologies; miRVana™ miRNA mimic, 4464066, ID MC10421; miRVana™ miRNA inhibitor, 4464084, ID MH10421) or miR-324-5p (10 nmol/L; miRVana™ miRNA mimic, 4464066, ID MC10253; miRVana™ miRNA inhibitor, 4464084, ID MH10253) or a combination of miR-181 and miR-324 together (10 nmol/L of each). After 5 hours, the media were changed and cells were exposed to hypoxia, or treated with TNF-α for 24-72 hours. Alternatively, on the following day, the untransfected and transfected cells were starved for 9 hours before caspase 3/7 assay. Transfection of cells with 20 pmol of Silencer Select negative control #1 (Ambion, 4390843), Silencer Select ETS-1 siRNA (Ambion, 432420, ID: S4847) and Silencer Select Notch4 siRNA (Ambion, 4392420, ID: S9643) was carried out with Lipofectamine RNAiMAX, as described above. The cells were used for experiments 24 hours post-transfection.

### Exosomal miRNA transfer under flow

6 × 10^3^ cells in 100 µL ECGM2 medium were seeded into each of six chambers of µ-Slide VI 0.4 (Ibidi, 80606). On the following day, the cells grown in the first chamber were transfected with fluorescent Cy3-miR (10 nmol/L, Ambion, AM17120) and washed several times before connecting the chamber to the other two flow chambers in tandem, so that the medium from chamber 1 would flow through chamber 2, then chamber 3 and out. A peristaltic pump (Ismatec REGLO ICC Digital Peristaltic Pump, Cole-Parmer, UK) created laminar flow of media over the endothelial cells at 4 dynes/cm^2^.

### Caspase 3/7 apoptosis assay

After an overnight incubation the cells were left in full medium or were incubated in serum- and growth factor-depleted medium for 9-24 hours to induce apoptosis. Apoptosis was measured using Cell Meter™ Caspase 3/7 Activity Apoptosis Assay Kit (AAT Bioquest, ABD-22796). Fluorescence intensity was analysed in Glomax™ luminometer at Ex/Em = 490/525 nm.

### NFκB luciferase reporter assay

NFκB activity was measured in luciferase reporter assay (Wojciak-Stothard et al., 2014a) in the Glomax™ luminometer.

### RNA Extraction

RNA was extracted from trypsinised HPAECs or lung tissue (∼10 mg) using miRCURY™ RNA Isolation Kit (Exiqon), according to manufacturer’s instructions.

### Real-time quantitative PCR

Input RNA (50-100 ng/µL) was reverse-transcribed using SuperScrip™ II Reverse Transcriptase (Invitrogen) or TaqMan MicroRNA Reverse Transcription Kit (Thermo Fisher Scientific) and with custom Multiplex RT Primer pool in a SimpliAmp™ Thermal Cycler (Applied Biosystems), according to the manufacturer’s instructions.

TaqMan^®^ Gene Expression Assays for KLF2 (Hs00360439_g1, Mm00500486_g1), Notch4 (Hs00965895_g1), ETS-1 (Hs00428293_m1), and GAPDH (Hs02786624_g1, Mm99999915_g1), and TaqMan^®^ miRNA Assays for miR-181a-5p (Assay ID 000480), miR- 324-5p (Assay ID 000539), and U6 snRNA (Assay ID 001973, all Thermo Fisher Scientific), were used to perform quantitative PCR (qPCR).

### Stimulation of KLF2 expression by shear stress

HPAECs were cultured in 9-cm^2^ Nunc^®^ Lab-Tek™ Flaskettes^®^ (VWR, 62407-340). The slides were detached and inserted into flow chambers in a parallel chamber flow apparatus (Wojciak-Stothard & Ridley, 2003). Endothelial cells were subjected to laminar flow at 10 dynes/cm^2^ for 24 hours. KLF2 and miRNA expression levels in cells grown under the static and flow conditions were measured by qPCR.

### EdU Proliferation Assay

The EdU Cell Proliferation Assay Kit (EdU-594, EMD Millipore Corp, USA, 17-10527) was used to measure the proliferation rate in cells, according to the manufacturer’s protocol. HPASMCs or HPAECs grown in serum-reduced, growth factor-depleted medium were stimulated with 10 ng/mL PDGF (Thermo Fisher Scientific, PHG0044) or 50 ng/mL of VEGF_165_ (R&D, 293-VE) 18 hours before the assay. Proliferation of ECFCs from healthy individuals and IPAH patients was evaluated in serum-reduced, growth factor-depleted media.

### RNA-Sequencing

Next-generation RNA-sequencing of HPAECs transfected with miR-181 or miR-324 was performed in duplicate at the Imperial BRC Genomics Facility (Imperial College of London, UK). RNA libraries were prepared using TruSeq^®^ Stranded mRNA HT Sample Prep Kit (Illumina Inc., USA) according to the manufacturer’s protocol. Libraries were run over 4 lanes (2 × 100 bp) on a HiSeq 2500 (Illumina Inc.) resulting in an average of 34.4 million reads per sample.

Sequence data was de-multiplexed using bcl2fastq2 Conversion Software v2.18 (Illumina Inc.) and quality analysed using FastQC. Transcripts from paired-end stranded RNA-Seq data were quantified with Salmon v0.8.2 using hg38 reference transcripts (Patro, Duggal et al., 2017). Count data was normalised to accommodate known batch effects and library size using DESeq2 (Anders, Pyl et al., 2015, Love, Huber et al., 2014). Pairwise differential expression analysis was performed based on a model using the negative binomial distribution and p-values were adjusted for multiple test correction using the Benjamini-Hochberg procedure. Genes were considered differentially expressed if the adjusted p-value was greater than 0.05 and there was at least a 1.5 fold change in expression. miRNA target prediction was carried out with TargetScan Human, miRecords and Ingenuity Expert Findings. Gene enrichment was carried out using Panther v14.0 (Thomas, Campbell et al., 2003).

### Animal experiments

All studies were conducted in accordance with UK Home Office Animals (Scientific Procedures) Act 1986. All animals were randomly allocated to groups, and all personnel involved in data collection and analysis (haemodynamics and histopathologic measurements) were blinded to the treatment status of each animal. Only weight-and age-matched males were included for experimentation as, in contrast to the human clinical studies, most animal studies have shown that female sex and estrogen supplementation have a protective effect against PAH(de Jesus Perez, 2011).

8-12 weeks old C57/BL male mice (20 g; Charles River, UK) were injected subcutaneously with Sugen (SU5416; 20mg/kg), suspended in 0.5% [w/v] carboxymethylcellulose sodium, 0.9% [w/v] sodium chloride, 0.4% [v/v] polysorbate 80, 0.9% [v/v] benzyl alcohol in deionized water once/week. Control mice received only vehicle. Mice were either housed in normal air or placed in a normobaric hypoxic chamber (10% O_2_) for 3 weeks (n= 8/group).

miR-181a-5p (ID MC10421) and miR-324-5p (ID MC10253) together, or Negative Control 1 mirVana™ miRNA mimics In Vivo Ready (Ambion Life Technologies) were complexed with Invivofectamine^®^ 3.0 reagent (Invitrogen) and injected intravenously twice a week at 2 mg per kg body weight. At 3 weeks, the mice were anaesthetised by intraperitoneal injection of Ketamine/Dormitor (75 mg/kg + 1 mg/kg). Development of PH was verified by the measurement of right ventricular systolic pressure (RVSP), right ventricular hypertrophy assessed as the right ventricle to left ventricle/septum ratio (RV/LV+S) and muscularisation of small intrapulmonary arteries, as described previously (Wojciak-Stothard et al., 2014a).

### Immunostaining

Immunostaining of paraffin embedded lung sections was carried out as previously described (Wojciak-Stothard et al., 2014a).

### Western blot analysis

Primary antibodies used in western blot analysis included goat polyclonal anti-KLF2 antibody (1:1000; Novus Biologicals, NB100-1051), mouse monoclonal anti-β-actin (1:3000; Sigma-Aldrich, A2228), Notch4 (1:1000; Santa Cruz Biotechnology, sc-393893), ETS-1 (1:1000; Santa Cruz Biotechnology, sc-55581), α-SMA (1:1000; Sigma, A5228) and rabbit polyclonal anti-PCNA (1:500, Santa Cruz Biotechnology, sc-7907). Secondary antibodies were HRP-linked Goat Anti-Rabbit IgG (1:3000; Sigma-Aldrich, A6154) and HRP-linked Sheep Anti-Mouse IgG (1:2000; GE Healthcare Life Sciences, NAG31V). For the analysis of p65 NFκB localisation and phosphorylation, human/mouse p65 (1:1000, Santa Cruz Biotechnology, SC-372-G) and human/mouse pSer536-p65 (1:1000, Cell Signaling, 3033) were used (Pradere, Hernandez et al., 2016).

### Nuclear translocation of p65NFκB

Nuclear translocation of p65 NFκB was studied with Image J by measuring colocalisation of nuclear stain DAPI with p65NFκB stained with rabbit p65 antibody (1:1000, Santa Cruz Biotechnology, SC-372-G) and TRITC-labelled goat anti-rabbit antibody (1:200, Jackson ImmunoResearch Laboratories, 111-025-144) in confocal images of cells or lung sections. The white pixel area, marking nuclear NFκB, was used to quantitate p65 NFκB translocation in cells and tissues.

### RNAscope^®^ *in situ* hybridisation

Histological sections of lung tissues from treatment-naïve PAH patients at lung transplantation (n=6), and control tissues comprising uninvolved regions of lobectomy specimens from 4 patients undergoing surgery for bronchial carcinoma and 2 unused donor lungs were from the tissue archives (Abdul-Salam, Wharton et al., 2010) at Hammersmith Hospital, Imperial College London, UK. Samples were fixed in 10% formal-saline and embedded in wax, and sections were processed for immunohistochemistry as previously described (Wojciak-Stothard, Abdul-Salam et al., 2014b). Sections from HPAH patients with c-terminal missense mutation in *KLF2* gene were obtained from 3 family members, father who died aged 32 and his two daughters who underwent lung transplantation (Eichstaedt et al., 2017). Lung sections from healthy and Sugen/hypoxia mice were also used for the analysis.

For formalin-fixed, paraffin-embedded tissues FFPE tissues, RNAscope^®^ Multiplex Fluorescent Reagent Kit v2 (Advanced Cell Diagnostics) and TSA™ Cyanine 3 & 5, TMR, Fluorescein Evaluation Kit System (PerkinElmer) were used according to manufactures’ protocols (Wang et al., 2012). Hybridization was carried out with target probes (human: Hs-ETS1-C1, NM_001143820.1; Hs-Notch4-C2, NM_004557.3; Hs-KLF2-C3, NM_016270.2; mouse: Mm-ETS1-C1, NM_011808.2; Mm-Notch4-C2, NM_010929.2; Mm-KLF2-C3, NM_008452.2).

### Angiogenesis assay

ECFC angiogenesis was studied in matrigel tube formation assay (Wojciak-Stothard et al., 2014a).

### Statistics

All experiments were performed at least in triplicate. Data are presented as means±SEM. Normality of data distribution was assessed with Shapiro-Wilk test in GraphPad Prism 7.03. Comparisons between 2 groups were made with Student t test or Mann-Whitney’s U test, whereas ≥3 groups were compared by use of ANOVA with Tukey’s post hoc test or Kruskal-Wallis with Dunn’s post hoc test, as appropriate. Statistical significance was accepted at P<0.05.

### Study approval

Venous blood samples were obtained with local ethics committee approval and informed written consent from healthy volunteers and idiopathic PAH patients. Participants were identified by number. Animal studies were conducted in accordance with UK Home Office Animals (Scientific Procedures) Act 1986.

## ACKNOWLEDGEMENTS

This research was supported by PhD Studentships from the Government of Saudi Arabia (Hebah Sindi) and British Heart Foundation project grant PG/16/4/31849.

We thank the staff of the Imperial NIHR/Imperial Clinical Research Facility, Hammersmith Hospital (London UK), and Dr John Wharton for their help in acquiring cells and lung sections from IPAH patients. We acknowledge the support from the Netherlands CardioVascular Research Initiative; the Dutch Heart Foundation, Dutch Federation of University Medical Centres, the Netherlands Organisation for Health Research and Development and the Royal Netherlands Academy of Sciences.

## AUTHOR CONTRIBUTIONS

HS, MA, AJA performed *in vitro* experiments, analysed data; KBJ, GR and VBA-S *in vitro* and *in vivo* experiments, immunohistochemistry, analysed data; BQC exosome quantification; RS and HJB provided ECFCs and critical analysed the manuscript; CCM analysed RNA-Seq data; SP, CAE and EG provided HPAH materials and analysis; CR and MR provided IPAH patient microRNA data and critically evaluated the manuscript; BWS secured funding, performed experiments, wrote the manuscript.

